# Chromosomal Inversion Symmetry: Generalized Chargaff Rules

**DOI:** 10.1101/013789

**Authors:** Sagi Shporer, Benny Chor, David Horn

**Affiliations:** School of Computer Science, Tel Aviv University, Tel Aviv 69978, Israel; School of Physics and Astronomy, Tel Aviv University, Tel Aviv 69978, Israel

**Keywords:** generalized Chargaff rules, chromosome k-mer distributions

## Abstract

The generalization of the second Chargaff rule to values of k larger than 1, states that the frequency of any k-mer on a single strand almost equals that of its inverse (reverse-complement). We demonstrate the validity of the generalized rule up to k=10 for all human chromosomes. Moreover, this **Inversion Symmetry** holds for many species, both eukaryotes and prokaryotes, for ranges of k which may vary from 7 to 10 as chromosomal lengths vary from 2Mbp up to 200 Mbp. We demonstrate that the statistical distributions of inverted pairs of k-mers are very different from other natral pairings of k-mers, implying that inversion symmetry is a basic principle of chromosomal structure. We suggest that it came into being because genomic evolution employed many rearrangements which consisted of inversions of chromosomal sections; on length scales down to order 1-10Kbp. Model simulations substantiate this claim.Low-scale inversions during chromosomal evolution imply that IS may exist for short sections of human chromosomes. This is indeed the case: we find that chromosome sections of length 5Kbp satisfy IS for k=1 and k=2. The largest value of k for which IS holds,which we call the k-limit of IS, increases logarithmically as the section length increases. The logarithmic dependence of the k-limit on the length of the chromosome is a universal characteristic, observed throughout the tree of life.

## Introduction

Erwin Chargaff has stated, in 1950, the important observation that the numbers of nucleotides in DNA satisfy #A=#T and #G=#C (Chargaff 1950, 1951). This statement, made on the basis of experimental observations with fairly low accuracy, played a crucial role in realizing that DNA has an underlying base-pair grouping, as proposed subsequently by Crick and Watson (1953) in their double-helix structure.

The second Chargaff rule (Rudner et al. 1968) states that the same sets of identities of nucleotide pairs hold for each long enough single DNA strand. This rule has been tested by (Mitchell and Bridge 2006) for genome assemblies of many species, and found to be globally valid for eukaryotic chromosomes, as well as for bacterial and archaeal chromosomes. It fails for mitochondria, plasmids, single-stranded DNA viruses and RNA viruses.

The validity of the second Chargaff rule is unexpected. Obviously it should be regarded as a global rule, i.e. applicable to large sections of chromosomes. Nonetheless, not being derived from a compelling principle such as the one underlying the first rule, it remains a mystery. This is even more so, when one studies extended versions of Chargaff’s second rule. Indeed (Albrecht-Buehler et al. 2006) observed that for triplet oligonucleotides, or 3-mers, it remains true that their chromsome-wide frequencies are equal to those of their reverse-complement 3-mers. Prabhu (1993) has shown that this symmetry holds up to 5-mers in various species. This has been reviewed by (Baldi and Brunak 2001) who have argued that such symmetry rules have to be incorporated in Markov models of genomic sequences.

We refer to the symmetry between numbers of appearances of k-mers and their reverse complements as

> **Inversion Symmetry (IS):** the number of occurrences of a k-mer of nucleotides on a chromosomal strand is almoste qual to that of its inverse (reverse-complement) string.

Note that this implies that the number of times a string of nucleotides of length *k* is observed on a strand, when read from 5’ to 3’, is almost equal to the number of times it is observed on the other strand when the latter is read from its 5’ end to 3’ end. Suggesting a criterion for exactness of IS by requiring that inequalities between frequencies of inverted k-mer pairs be less than 10%, we will show that the IS is valid up to k=10 on long human chromosomes. We will refer to the highest k for which IS is valid as the k-limit of inversion symmetry.

By comparing inverted pairs with other natural pairings of k-mers, we will demonstrate the unique features of IS, separating it from other pairings. Moreover, we will argue that IS should not be regarded just as a feature to be imposed on chromosome modeling, but also as one reflecting evolutionary dynamics of chromosomes. We will demonstrate that in models invoking random inversions of chromosome sections, one can obtain IS k-limits that mimic the biological ones. The values of k-limits, both the ones observed in different species and the ones derived from models, increase logarithmically with chromosome length.

We will also discuss CpG effects on the distributions of other k-mer pairings, and the fact that IS exists for both unmasked and masked version of chromosomes, demonstrating that it is not due to repetitive and low-complexity sequences.

## Results

### Inversion symmetry (Generalized Chargaff Rule)

Let S and S^*^ be two strings of nucleotides of same length k. Suppose they appear N(S) and N(S^*^) times respectively on a chromosome. We denote by x(S,S^*^) the relative difference x(S,S^*^)=|N(S)-N(S^*^)|/(N(S)+N(S^*^). In the following we will look at values of the variable x(S,S^*^) over all possible choices of inverse pairs, and demonstrate that they are distributed differently than other types of k-mer pairs. Moreover, we will evaluate the average values, E_k_[x], and use them to demonstrate and quantify IS.

Let us start with the latter, computing E_k_[x] for different k on various chromosomes of the most recent human assembly, HG38. Data were downloaded from the UCSC genome browser https://genome.uscs.edu. The calculated values of E_k_[x] for several human chromosomes are displayed as function of k in Fig. 1. Inversion Symmetry (IS) is seen to hold quite well for k-mers with high k-values for all the displayed chromosomes. Chr Y, which is the shortest among the 24 chromosomes, has the worse inversion symmetry. IS holds also for all other (not shown) chromosomes but fails (even at the k=1 level) for the mitochondrial one.

**Fig. 1.**
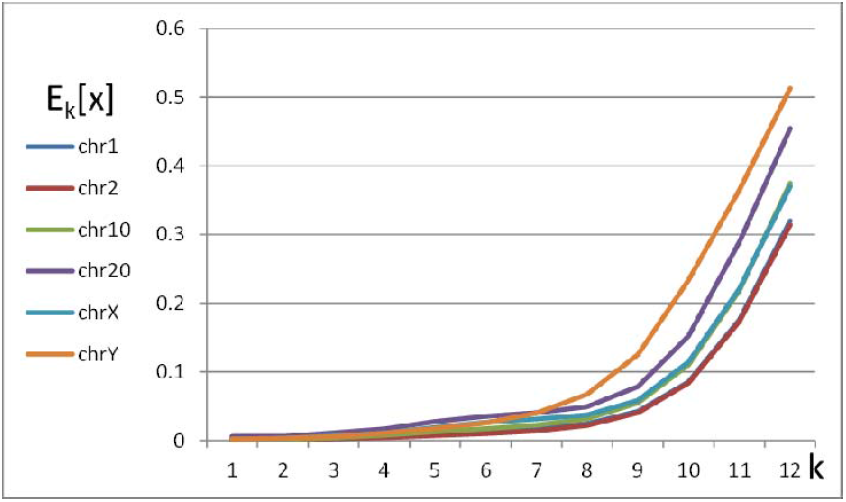
Averages of relative differencess between occurrences of k-mers and their inverses (reverse-complements), E_k_[x], for different chromsomes of the HG38 human assembly, plotted *vs* k.

Repetitive structures are well-known to constitute major fractions of eukaryotic chromosomes, hence one may wonder to what extent they are responsible for the observed inversion symmetry. To resolve this issue, we employed the same operations on the masked output of the UCSC genome browser, after screening chromosomes for interspersed repeats and low complexity sequences. The results (see Supplemental Material) keep displaying the same behavior, with negligible differences for high values of k. Even ChrY, which is the most notorious hub of repeats, with only 36% of it surviving the masking filter, keeps showing the same qualitative behavior as in Fig. 1. In the Supplemental Material (Table S1) we provide a list of the highest k-values for which E_k_[x]<0.1, which we call the k-limits of IS, both before and after masking. The observed reduction in k-limits from 10 to 9 for the largest chromsomes, may well be just because filtering shortens the effective chromsome lengths. The effect of length on k-limits is an issue to which we will return below.

We have performed the same analysis on the older genome assembly HG18, leading to very similar results (see Supplemental Material Table S2). We find similar IS results for mouse, frog, fly, worm, and yeast. Moreover, we find that inversion symmetry holds also for bacteria, but it is valid for a lower range of k-mers, only up to k=6 or 7.

### Outstanding features of inverted k-mer pairs

In order to demonstrate how Inversion Symmetry, observed for frequencies of inverted pairs, differs from other natural pairings, we compare here three different choices of pairings of k-mers,

a. - Inverted pairs (e.g. CGA *vs* TCG)
b. - Random pairs
c. - Reversed pairs (e.g. CGA *vs* AGC)

For all three types of pairings we will draw histograms of x(S,S^*^b^*^)=|N(S)- N(S^*^)|/(N(S)+N(S^*^), and evaluate their averages, E_k_[x(S,S^*^)].

Fig. 2a depicts the distribution of inverted pairs on human chr 1 of HG38, evaluated for k=4 to 10. These distributions are very narrow, befitting very low E_k_[x] values, of the type displayed in Fig. 1. As k increases they widen, leading to increasing average E_k_[x] values, which will be discussed below and quoted in Table 1. In Figs. 2b and 2c we plot the corresponding distributions for the cases of random pairs (b) and reversed pairs (c) on chr 1. Note that these distributions are completely different: they possess a rugged wavy behavior, stretching over the whole range of 0<x<1. Similar distributions are also observed to occur for masked chromsomes.

**Fig. 2.**
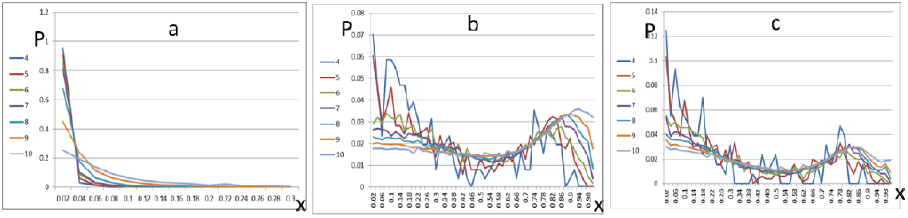
HG38 chrl: Histogram (probability distribution in bins of Δx=0.02) of relative occurrences of k-mer pairs vs x for different values of k (4 to 10). a: inverted pairs; plotted range is x<0.3, below which the histogram values are negligibly small. b: random pairs for full x range; c: reversed pairs for full x range.

**Table 1.** comparisons of averages E_k_[x] of μ_ka_=inverted pairs, μ_kb_=random pairs, and μ_kc_=reversed pairs, for chrl of HG38.

| **k** | **$\mu_{ka}$** | **$\mu_{kb}$** | **$\mu_{kc}$** | **$\mu_{ka}/\mu_{kb}$** | **$\mu_{ka}/\mu_{kc}$** |
| --- | --- | --- | --- | --- | --- |
| 1 | 0.0009 | 0.0828 | 0.0000 | 0.0114 |  |
| 2 | 0.0008 | 0.1993 | 0.1547 | 0.0042 | 0.0054 |
| 3 | 0.0031 | 0.2619 | 0.2060 | 0.0119 | 0.0151 |
| 4 | 0.0055 | 0.3278 | 0.2665 | 0.0169 | 0.0208 |
| 5 | 0.0090 | 0.4048 | 0.3154 | 0.0223 | 0.0286 |
| 6 | 0.0126 | 0.4392 | 0.3623 | 0.0288 | 0.0349 |
| 7 | 0.0171 | 0.4864 | 0.4008 | 0.0352 | 0.0427 |
| 8 | 0.0247 | 0.5191 | 0.4333 | 0.0475 | 0.0570 |
| 9 | 0.0426 | 0.5465 | 0.4607 | 0.0780 | 0.0925 |
| 10 | 0.0853 | 0.5707 | 0.4883 | 0.1494 | 0.1746 |
| 11 | 0.1758 | 0.6029 | 0.5283 | 0.2917 | 0.3328 |
| 12 | 0.3199 | 0.6666 | 0.6034 | 0.4799 | 0.5302 |

The vast difference between case (a) and cases (b) and (c) should be kept in mind when we summarize the observations in terms of only the averages, μ_k_=E_k_[x], in all three cases, to be denoted by μ_ka_, μ_kb_ and μ_kc_. They are presented in Table 1. We note that the values of μ_ka_, μ_ka_/μ _kb_ and μ_ka_/μ_kc_ keep increasing with k. Let us (quite arbitrarily) set the bounds

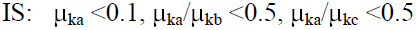

as defining the validity criteria of Inversion Symmetry. They are satisfied up to k=10 in the example of chr1 in Table 1. A condition like (μ_ka_ / (μ_kc_ <0.5 is meant as one indication of the difference between the two distributions, which differ by much more than their averages, as seen in Figs. 2a and 2c.

Similar results can be obtained for almost all species, both eukaryotes and prokaryotes. Examples are provided in the Supplemental Material. Here we display, in Table 2, the results for chr 4 of S. cerevisiae. Clearly these data allow for a lower range of up to k=7 using our criteria for IS validity. The distributions of the three types of k-mer pairings are displayed in Figs. 3a, 3b and 3c.

**Table 2.**
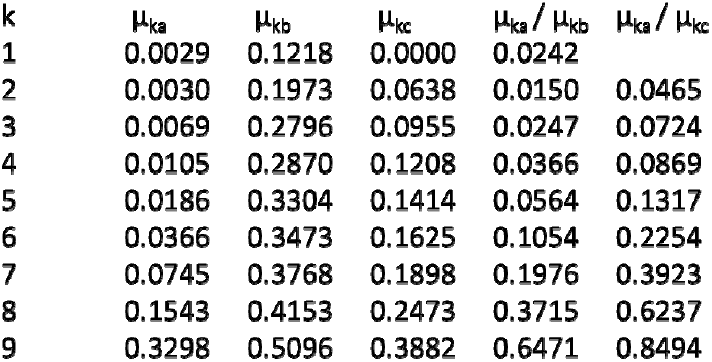
comparisons of averages E_k_[x] of μ_ka_=inverted pairs, μ_kb_=random pairs, and μ_kc_=reversed pairs, for chr 4 of S. cerevisiae.

| **k** | **$\mu_{ka}$** | **$\mu_{kb}$** | **$\mu_{kc}$** | **$\mu_{ka}/\mu_{kb}$** | **$\mu_{ka}/\mu_{kc}$** |
| --- | --- | --- | --- | --- | --- |
| 1 | 0.0029 | 0.1218 | 0.0000 | 0.0242 |  |
| 2 | 0.0030 | 0.1973 | 0.0638 | 0.0150 | 0.0465 |
| 3 | 0.0069 | 0.2796 | 0.0955 | 0.0247 | 0.0724 |
| 4 | 0.0105 | 0.2870 | 0.1208 | 0.0366 | 0.0869 |
| 5 | 0.0186 | 0.3304 | 0.1414 | 0.0564 | 0.1317 |
| 6 | 0.0366 | 0.3473 | 0.1625 | 0.1054 | 0.2254 |
| 7 | 0.0745 | 0.3768 | 0.1898 | 0.1976 | 0.3923 |
| 8 | 0.1543 | 0.4153 | 0.2473 | 0.3715 | 0.6237 |
| 9 | 0.3298 | 0.5096 | 0.3882 | 0.6471 | 0.8494 |

**Fig. 3.**
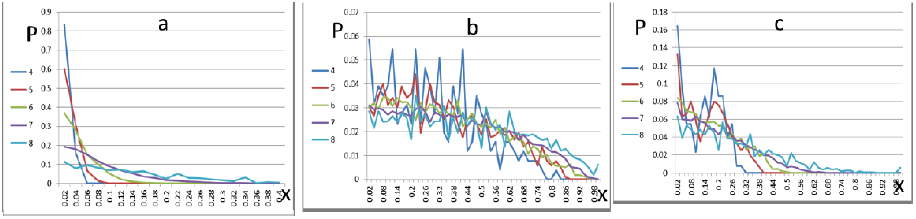
S. cerevisiae chr4: Histogram of relative occurrences of k-mer pairs vs x for different k. a: inverted pairs; range x<0.4. b:random pairs for full x range. C: reverse pairs for full x range.

One should realize that chr 4 of S. cerevisiae, used for this analysis, is of length 1.5Mbp, while the length of human chr1 in HG38 is 230Mbp. This difference by two orders of magnitude is part of the reason why the human chromosome displays a higher k-limit of inversion symmetry. One obvious effect of the length is the larger fraction out of all possible k-mers that can be realized within the measured strand. In chr 4 of S. cerevisiae, we find that for k=10 only 0.77 of all possible k-mers exist in the genomic sequence, and this number reduces to 0.40 and 0.14 for k=11 and 12. For such low coverages of all k-mers there are many cases where a string S appears while its inverse S^*^ does not, therefore increasing E_k_[x]. By comparison, in human chr 1 we find that for k=11 and 12, 0.99 and 0.91 of all possible k-mers exist in the data.

### The CpG effect

The large hump in the distribution of reverse-pairs and random pairs in human deserves some elaboration. This is related to the well-known CpG suppression in tetrapods, i.e. the very low number of appearances of CG compared to all other dimers on their genomes.

CpG suppression has a substantial effect on x-distributions of reversed pairs. Hence we have reanalyzed all paired distributions after eliminating all k-mers which carry a CG dimer. The results are displayed in Figs 4a to 4c. Since the CG dimer is the inverse of itself, it is no wonder that the distribution of all inverse pairs looks still the same even when all CG dimers are eliminated, as shown in Fig. 4a. However in the other cases, we see that by CG removal the hump at large x values was eliminated. This hump is therefore an effect of CpG suppression. In S. cerevisiae, where CpG is not suppressed, this hump is indeed absent (see Figs. 3b and 3c). One should note that, even in in Fig. 4, in spite of the hump removal, the distribution of inverted pairs is still much narrower than for the other k-mer pairings.

**Fig. 4.**
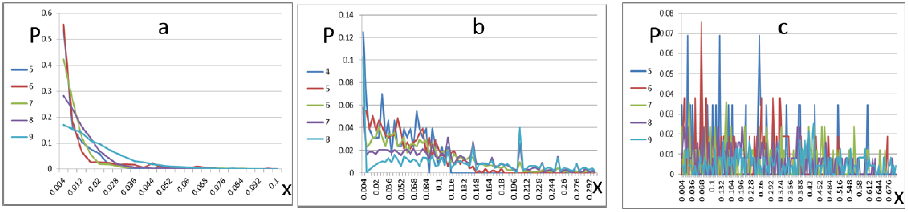
HG38 chr 1, after elimination of all k-mers containing a CG dimer: Histogram of relative occurrences of k-mer pairs vs x for different k. a: inverted pairs; range x<0.1. b: random pairs; range x<0.7. c: reversed pairs; range x<0.6.

### Modeling Inversion Symmetry

If IS holds for some k=k_0_, it will hold also for all k<k_0_, since the latter are substrings of the former and, therefore, all the frequencies of the k inverted-pair substrings will be matched since the frequencies of their k_0_ hosts are being matched. One may wonder to what extent the opposite may hold within, e.g., low order Markov models: will a Markov model, constructed such that it satisfies IS for some k account for IS at the level k+1? The answer is negative. Even for low k values, a Markov model based on a lower statistic cannot generate the higher statistic (Baldi and Brunak 2001).

The simplest random model is that of a uniform distribution, which is generated on the basis of the second Chargaff rule (i.e. #A=#T and different from #C=#G). Such a distribution will trivially account for low (μ_ka_ values for large values of k, limited by the length of the model chromsome. However it will also give rise to very low (μ_kc_ values for a similar range of k, because any comparison of k-mers with one of their permutations will lead to similar E_k_[x]. In other words, this random independent (but not IID) model satisfies additional symmetries that are not observed in genomic data. Therefore it cannot serve as a model of inversion symmetry.

A plausible explanation of the observed IS can be based on the fact that genomes evolve through rearrangement processes. By comparing synteny blocks in human and mouse, (Pevzner and Tesler 2003) have argued that rearrangements occur on many scales in the genome, and intra-chromosomal rearrangements are more frequent than inter-chromosomal ones. Rearrangements may be viewed as inversions of sections between two breakpoints on the chromosome, and they may even follow one another in a nested fashion. In their study (Pevzner and Tesler 2003) demonstrated that human and mouse chr X share 281 synteny blocks of size >1Mb, and at least 245 rearrangements occurred since the divergence of the two species.

Building on this intuition, derived from comparative genomics, we suggest that a series of such rearrangements on different scales may lead to IS. We demonstrate it on a simple model, starting from the human mitochondrial chromosome, which does not satisfy the second Chargaff rule. Since the mitochondrial chromosome is only 16Kbp long, we first construct out of it an enlarged model chromosome with length L= 100Mbp, by concatenating random selections of subsequences of chr M. We then apply to it rearrangements at various scales. We found that 5,000 rearrangements at scales of 100K have led to good IS effects, but best results were obtained for 50,000 rearrangements, whose breakpoints were randomly chosen, and their section lengths befit a uniform distribution of length <10K. These results exhibit a high degree of IS, as displayed in Fig. S1 of the SM.

Next we have also tested the application of random inversions to random models. A simple model of 1^st^ order statistics is not good enough, because multiple inversions may lead to symmetries higher than IS. Trying Markov models based on various random choices of transition probabilities among nucleotides one can obtain IS even if the original Markov chain does not possess any particular symmetry, if sufficiently many inversion rearrangements have been applied. Choosing sections of various lengths, with lengths uniformly distributed within a range R, for inversion processes, and applying such inversions for G generations, we find that for model chromosomes of L=1M and R=1K or 10K, we can obtain IS of k-limit=5 with G=10K, and k-limit=6 with G=100K. Increasing G to 1M already leads into the zone of large reversal symmetries. For L=10M and R=10K one induces IS up to k-limit of 7 with G=1M and 8 with G=8M.

### Inversion Symmetry for Chromosomal Sections

In view of the models discussed above one may expect IS to be observed on many sections of large chromosomes, as long as these sections are large enough so that they are expected to experience sufficiently many rearrangements during evolution. We have tested it on human genome assemblies. In Fig. S2 of the Supplemental material we display a characteristic distribution of inverted pairs drawn from a section of length 10Mbp, and in Fig. S3 we show an analogous distribution for length of 1Mbp. The IS quality, as determined by our convention, deteriorates leading to lower k-limits as the length of the section decreases, but it remains visible. The distributions in Fig. S3 are evidently noisier than their analogs in Fig. S2; however they are much narrower than those of the reversed and random pairs (not shown here).

To study systematically different sections of chromosomes, we evaluate the E_k_[x] values of inverted, random and reverse pairs, on non-overlapping windows of given lengths L. In practice, all inverted pairs lead to smaller results than the other pairing choices. To determine the k-limit we impose the E_k_[x] <0.1 on the average of all trials of inverted pairs. The example displayed in Fig. S4 is of chr1, which is being tested with windows of length L=5Kbp for inverted pairs of k=2. Although their average value is 0.07, obeying our criterion for IS validity, it is quite obvious that on many 5K windows their values are higher. The value k=2 is chosen as the k-limit of IS validity in this case. Reducing the section length further down to L=1Kbp, we find that IS fails even at order k=1, i.e. the second Chargaff rule does not hold.

Similar evaluations for different chromosomes, on both HG18 and HG38 assemblies, lead in a consistent manner to the k-limits of “human sections” displayed in Table 3, where they are compared with results obtained for various other species, both eukaryotes and prokaryotes. They all follow a logarithmic increase with the length of the chromosome, or chromosomal section, as is quite evident from their display in Fig. 5.

**Table 3.** Maximal k-values, establishing IS limits for human data as well as other eukaryotes and prokaryotes.

| species | length | k-limit |
| --- | --- | --- |
| HG38.chr1 | 230479627 | 10 |
| HG18.chr1 | 224999368 | 10 |
| chimp.panTro2.chr1 | 217189828 | 10 |
| mouse.mm10.chr1 | 191908761 | 10 |
| HG18.chrX | 151058618 | 9 |
| zebrafish.danRer6.chr7 | 76727960 | 9 |
| melanogaster.dm3.chr3R | 27905045 | 9 |
| elegans.ce10.chrV | 20924149 | 9 |
| HG18.chrY | 25652849 | 8 |
| human section of 10M | 10000000 | 8 |
| Escherichia_coli_K_12_substr__W3110 | 4646325 | 8 |
| Bacillus_subtilis_uid76 | 4215599 | 8 |
| human section of 5M | 5000000 | 7 |
| Mycobacterium_avium_paratuberculosis | 4829775 | 7 |
| Pyrococcus_furiosus_uid287 | 1908250 | 7 |
| Thermotoga_maritima_uid111 | 1860719 | 7 |
| cerevisiae.sacSer3.chrIV | 1531933 | 7 |
| human section of 1M | 1000000 | 6 |
| human section of 100K | 100000 | 5 |
| human section of 50K | 50000 | 4 |
| human section of 10K | 10000 | 3 |
| human section of 5K | 5000 | 2 |

**Fig.5.**
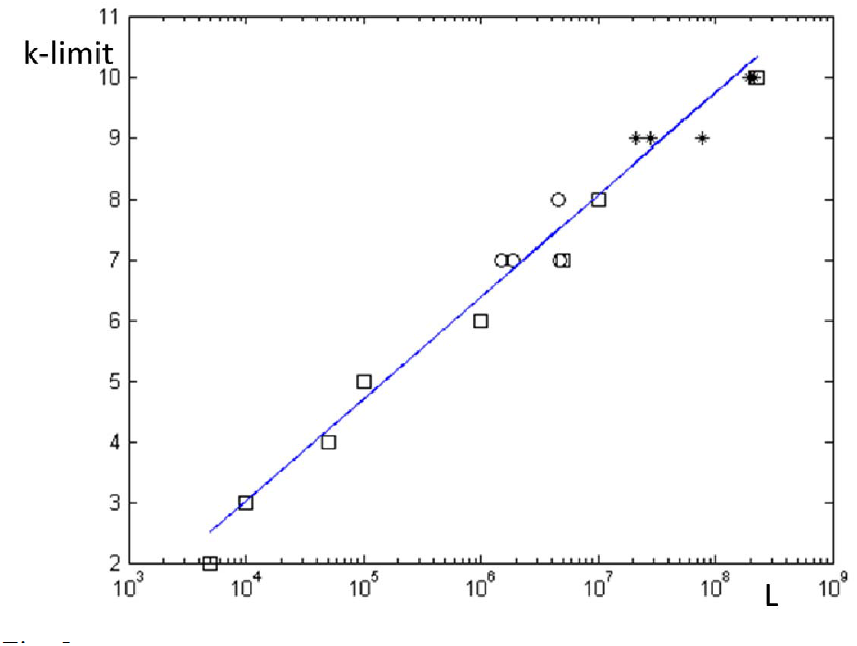
k-limits *vs* chromosomal length, taken from Table 3, display universal logarithmic behavior. Boxes are human data, stars denote other eukaryotes, and circles represent prokaryotes. The shown fit to this set of data is 0.73^*^log_10_(length), and should serve as an indication of the observed logarithmic increase of the k-limits.

## Discussion

Generalizing the second Chargaff rule to k-mers with 1<k≤10 or so, we have demonstrated the existence of an **Inversion Symmetry**, stating that the frequency of any particular k-mer is equal to that of its inverse (reverse-complement) on the same strand. This is tantamount to stating that the k-mers encountered on one strand, when read from 5’ to 3’, are the same as those encountered on the other strand when read from its 5’ end to 3’ end. Examining both eukaryotes and prokaryotes, we find that IS holds for a large range of k, which grows logarithmically with the length of the chromosome (or chromosome segment). We have introduced an IS criterion of μ_k_=E_k_[x]<0.1 for inverted pairs; moreover, comparing a:inverted pairs with b: random pairs and c:reversed pairs we have also demanded μ_ka_/μ_kb_<0.5,μ_ka_/μ_kc_<0.5, This defines what we mean by IS quantitatively, and has been applied to all cases that we have studied.

We have demonstrated that the statistics of inverted pairs on long chromosomal sections are very different from those of random pairs or reversed pairs. This indicates that IS must come about through active processes, which have shaped chromosomes into this large-scale behavior, found in genome assemblies. We have proposed that the mechanism for IS emergence is primarily due to chromosomal rearrangements throughout the evolutionary history of chromosomes. We demonstrated this effect on several synthetic models, starting with an “asymmetric chromosome” which violates IS even at k=1, and ending with explicit IS with high k-limits. According to the model, these rearrangements include many inversions of small genomic sections in order to produce IS for large k-values.

A glimpse at the ubiquity of inversions was recently provided by (Chaisson et al. 2014). Comparing their analysis of the haploid human genome CHM1 with the assembly of HG19 they have listed 14 high-confidence inversion calls (Supplementary Table 15 of their paper), involving one of size 1.09M, one of size 220K, one of 12K and the rest of few Kbp lengths. Observing such inversions on existing human data is very suggestive that they have been part and parcel of genomic evolution throughout evolutionary history.

Fig. 5 summarizes the universal behavior of k-limits of IS on chromosomes of both eukaryotes and prokaryotes. The major limiting factor is the length of the chromosome, or chromosomal section. Masked human chromosomes, with low complexity genomic sections removed (see Tables S1 and S2 of SM) fall also in line with this general behavior. Our model suggests that large numbers of small inversions are needed, in order to implement the creation of large numbers of instantiations of inverted pairs. We conclude therefore that large chromosomal lengths play important roles in both allowing for the appearance of all k-mer instantiations for high k, and for providing enough space so that many inversions lead to IS without introducing too large symmetries among pairings of k-mers which are permutations rather than inversions of one another. All this eventually leads to the logarithmic increase of k-limits with chromosomal length.

We found that generalized Chargaff rules hold, up to k=2, even for very short sections of human chromosomes, e.g. of size 5Kbp. An example of what happens when one tests the Chargaff rule on non-overlapping windows of size 1 Kbp is shown in Fig. S5. We see that the rule fails, but it also displays very non-homogenous behavior. This may be related to reports in the literature that there exists an excess of G+T over A+C on the coding strand, within most genes. Green et al. (2003) have argued that mutational asymmetry has acted over long periods of time to produce such a compositional asymmetry, and jumps of such asymmetries are associated with loci of replication origin. These questions have also been studied by Huvet et al. (2007).

Returning to the large scale picture, of chromosomes with lengths of 2Mbp to 200Mbp, we reiterate our main conclusions: inversion symmetry has been demonstrated to hold with k-limits varying from 7 to 10. Its accuracy is quite surprising, especially when compared with other pairings of k-mers. It is therefore important to understand its origin. We suggest it comes about through chromosomal rearrangements, which involved inversions at various length scales throughout the history of genomic evolution.

